# BEARscc determines robustness of single-cell clusters using simulated technical replicates

**DOI:** 10.1101/118919

**Authors:** DT Severson, RP Owen, MJ White, X Lu, B Schuster-Böckler

## Abstract

Technical variance is a major confounding factor in single-cell RNA sequencing, not least because it is not possible to replicate measurements on the same cell. We present BEARscc, a tool that uses RNA spike-in controls to simulate experiment-specific technical replicates. We demonstrate that the tool improves the unsupervised classification of cells and facilitates the biological interpretation of single-cell RNA-seq experiments.

## Introduction

Single cell messenger RNA sequencing (scRNA-seq) is a powerful tool to study cell subpopulations relevant to disease and development, including rare cell types^1,2^. However, scRNA-seq has inherently high technical variability and it is not possible to have true technical replicates for the same cell, presenting a major limitation for scRNA-seq analysis^3,4^. Specifically, read count measurements often vary considerably as a result of stochastic sampling effects, arising from the limited amount of starting material^3,4^. Also, false-negative observations frequently occur because expressed transcripts are not amplified during library preparation (the “drop-out” effect)^3,4^. Another common problem is systematic variation due to minute changes in sample processing. These batch-dependent differences in cDNA conversion, library preparation and sequencing depth can easily mask biological differences among cells and might compromise many published scRNA-seq results^5^.

One widely adopted approach to adjust for technical variation between samples is the addition of known quantities of RNA “spike-ins” to each cell sample before cDNA conversion and library preparation^6^. Several methods use spike-ins to normalize read counts per cell before further analysis^7,8^, but this use has been criticized because it exacerbates the effect of differences in RNA content per cell, e.g. due to variations in cell size^7,9^. Unfortunately, the limited volumes of starting material in single-cell transcriptomics inherently preclude the possibility of true technical replication.

To address this shortcoming of scRNA analysis, we developed “BEARscc”, an algorithm that uses spike-in measurements to model the distribution of experimental technical variation across samples to simulate realistic technical replicates. The simulated replicates can be used to quantitatively and qualitatively evaluate the effect of measurement variability and batch effects on analysis of any scRNA-seq experiment, facilitating biological interpretation. BEARscc represents a novel use for spike-in controls that is not subject to the same problems as per-sample normalization.

## Results

### Outline of BEARscc workflow

BEARscc consists of three steps (Figure 1): modelling technical variance based on spike-ins (Step 1); simulating technical replicates (Step 2); and clustering simulated replicates (Step 3). In Step 1, an experiment-specific model of technical variability (“noise”) is estimated using observed spike-in read counts. This model consists of two parts. In the first part, expression-dependent variance is approximated by fitting read counts of each spike-in across cells to a mixture model (see Methods). The second part, addresses drop-out effects. Based on the observed drop-out rate for spike-ins of a given concentration, the ‘*drop-out injection distribution*’ models the likelihood that a given transcript concentration will result in a drop-out. The ‘*drop-out recovery distribution*’ is estimated from the drop-out injection distribution using Bayes’ theorem and models the likelihood that a transcript that had no observed counts in a cell was a false negative.

**Figure 1.**
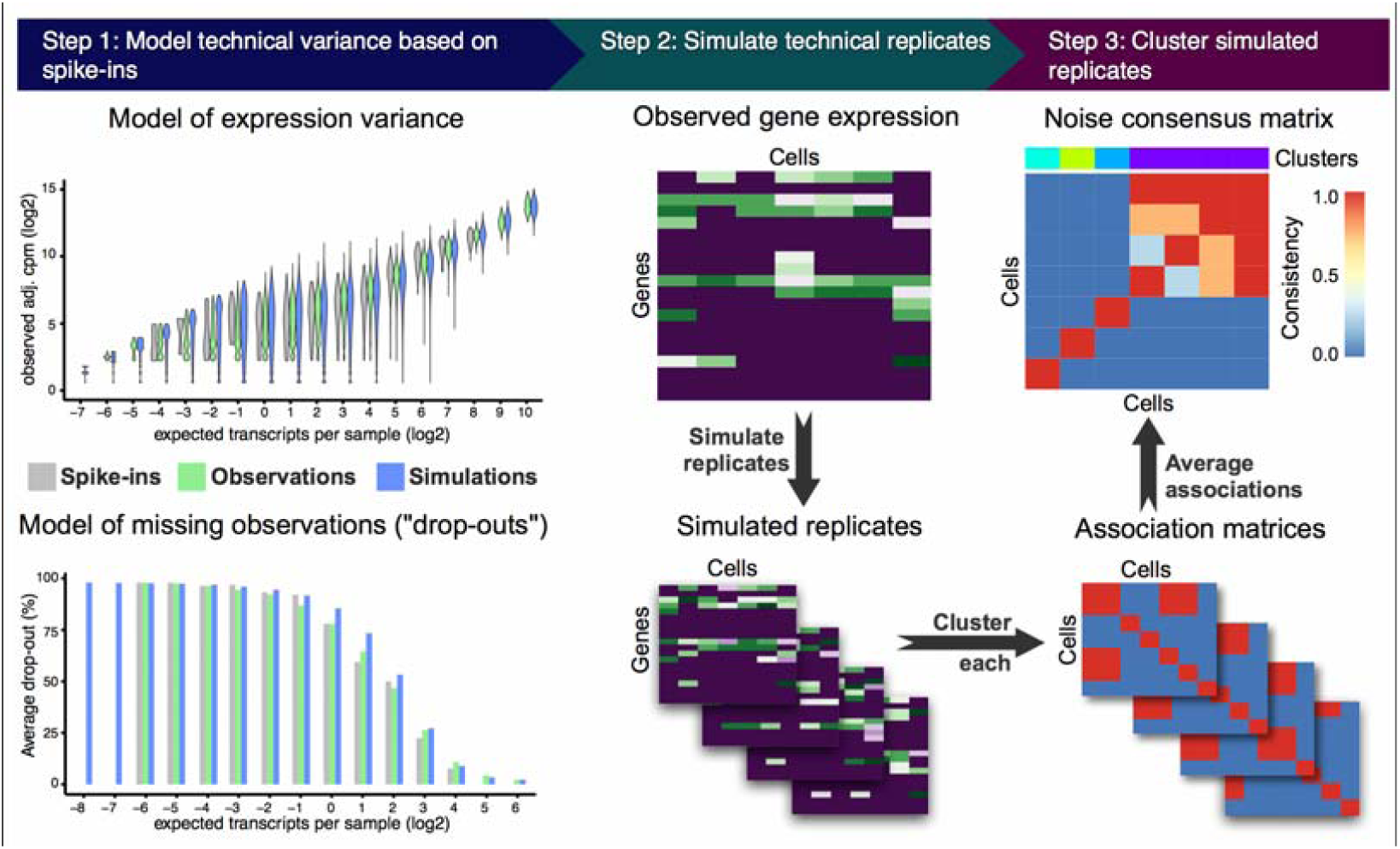
Overview of the BEARscc algorithm. **Step 1**, The variance of gene expression expected in a replicate experiment is estimated from the variation of spike-in measurements. Top: variation in spike-in read counts corresponds well with experimentally observed variability in biological transcripts (for details of control experiment see Methods) and read counts simulated by BEARscc. Bottom: Drop-out likelihood is modelled separately, based on the drop-out rate for spike-ins of a given concentration. Shown is the average percentage drop-out rate as a function of the number of transcripts per sample, for spike-ins, simulated replicates and experimental observations in a control experiment (see Methods). **Step 2**, simulating technical replicates: the observed gene counts (top matrix) are transformed into multiple simulated technical replicates (bottom) by repeatedly applying the noise model derived in Step 1 to every cell in the matrix. **Step 3**, calculating a consensus: simulated replicate (from Step 2) is clustered create an association matrix. All the association matrices (bottom) are averaged into a single *noise consensus matrix* (top) that reflects the frequency with which cells are observed in the same cluster across all simulated replicates. Based on this matrix, noise consensus clusters can then be derived (colored bar above matrix).

In Step 2, BEARscc applies the model from Step 1 to produce simulated technical replicates. For every observed gene count below which drop-outs occurred amongst the spike-ins, BEARscc assesses whether to convert the count to zero (using the drop-out injection distribution). For observations where the count is zero, the drop-out recovery distribution is used to estimate a new value, based on the overall drop-out frequency for that gene. After this drop-out processing, all non-zero counts are substituted with a value generated by the model of expression variance (from Step 1), parameterized to the observed counts for each gene. Step 2 can be repeated any number of times to generate a collection of simulated technical replicates.

Re-analyzing the simulated technical replicates in the same way as the original observations can reveal the robustness of the results to the modelled technical variation. Specifically, in Step 3 we focus on clustering analysis. Each simulated technical replicate is clustered using the same algorithm parameters as for the original observation. An association matrix is created in which each element indicates whether two cells share a cluster identity (1) or cluster apart from each other (0) in a particular replicate (Figure 1, step 3). We provide a visual representation of the clustering variation on a cell-by-cell level by combining association matrices to form the ‘*noise consensus matrix*’. Each element of this matrix represents the fraction of simulated technical replicates in which two cells cluster together (the ‘*association frequency*’), after using a chosen clustering method. To quantitatively evaluate the results, three metrics are calculated from the noise consensus matrix: ‘*stability*’ is the average frequency with which cells within a cluster associate with each other across simulated replicates; ‘*promiscuity*’ measures the association frequency between cells within a cluster and those outside of it; and ‘*score*’ is the difference between ‘*stability*’ and ‘*promiscuity*’. Importantly, ‘*score*’ reflects the overall “robustness” of a cluster to technical variance.

The ‘*score*’ statistic can also be used to address the challenge in cell type classification of determining the optimal number of clusters, *k*, into which cells are placed. Heuristics, such as the silhouette index or the gap statistic^10,11^, are commonly used but fail to account for expected variance in the similarity between cells. BEARscc's ‘*score*’ statistic provides an alternative approach to address this issue. Performing hierarchical clustering on the ‘noise consensus matrix’ allows BEARscc to split cells into any number of clusters between 1 and *N* (the total number of cells). The clustering with a maximum ‘*score*’ (within a biologically reasonable range) represents the optimal trade-off between within-cluster stability and between-cluster variability (see Methods).

### Evaluation with experimental replicates

To test the accuracy of BEARscc, we diluted one RNA-seq library derived from bulk human brain tissue to single cell RNA concentrations and sequenced 48 of these samples with ERCC spike-ins^12^; these are ‘real’ technical replicates to compare to the simulated technical replicates generated by BEARscc. The mean and variance of the simulated counts produced by BEARscc closely matched the experimentally determined values (Figure 1, step 1 - top; Supplementary Figure 1a,b). For 95% of the genes expressed in the library, the simulated drop-out rate differed from the observed drop-out rate by less than 9% (Figure 1, step 1 – bottom; Supplementary Figure 1c). Together, these results suggest that technical variation simulated by BEARscc closely resembles technical variation observed experimentally. The simulated expression of genes with less than 1 observed count deviated slightly from the experimentally determined values (Supplementary Figure 1a), however such small expression differences are unlikely to be reproducible as they fall outside the dynamic range of any single cell experiment.

To benchmark BEARscc, we performed a control experiment in which we sequenced 45 “blank” samples alongside the diluted brain RNA samples, in two batches. The blanks only contained spike-ins and trace amounts of environmental contamination, producing sporadic read counts. We clustered the data from the brain samples and blanks using three widely used clustering algorithms (RaceID2^13^, BackSPIN^14^, and SC3^15^), either alone or after simulating technical replicates using BEARscc. Correct clustering should give perfect separation of brain and blank samples. To avoid artifacts due to differences in amplification-dependent library size, we applied an adjusted cpm normalization^5^. Otherwise, standard parameters were used for all three clustering algorithms. As an alternative to BEARscc, we also tested a simple sampling approach where we repeatedly sampled half of all expressed genes and re-clustered the cells based on this subset (see Methods). Without BEARscc or this sampling approach, all three clustering algorithms created false-positive clusters (Figure 2a, Supplementary Figure 2a-c, top). BEARscc provided a clear improvement over the original clustering and the sampling approach (Figure 2a). Overall, BEARscc separated brain tissue and blank samples correctly and eliminated spurious clusters that corresponded to batch effects (Supplementary Figure 2a,c, colored bars above matrices). In the case of using BEARscc with RaceID2, three outlier cells were incorrectly identified to be “robust” clusters (Supplementary Figure 2b, colored bars above matrix); the libraries for these three samples contained fewer than 1,000 observed transcripts, indicating that BEARscc is limited by RaceID2's oversensitivity to library size differences.

**Figure 2.**
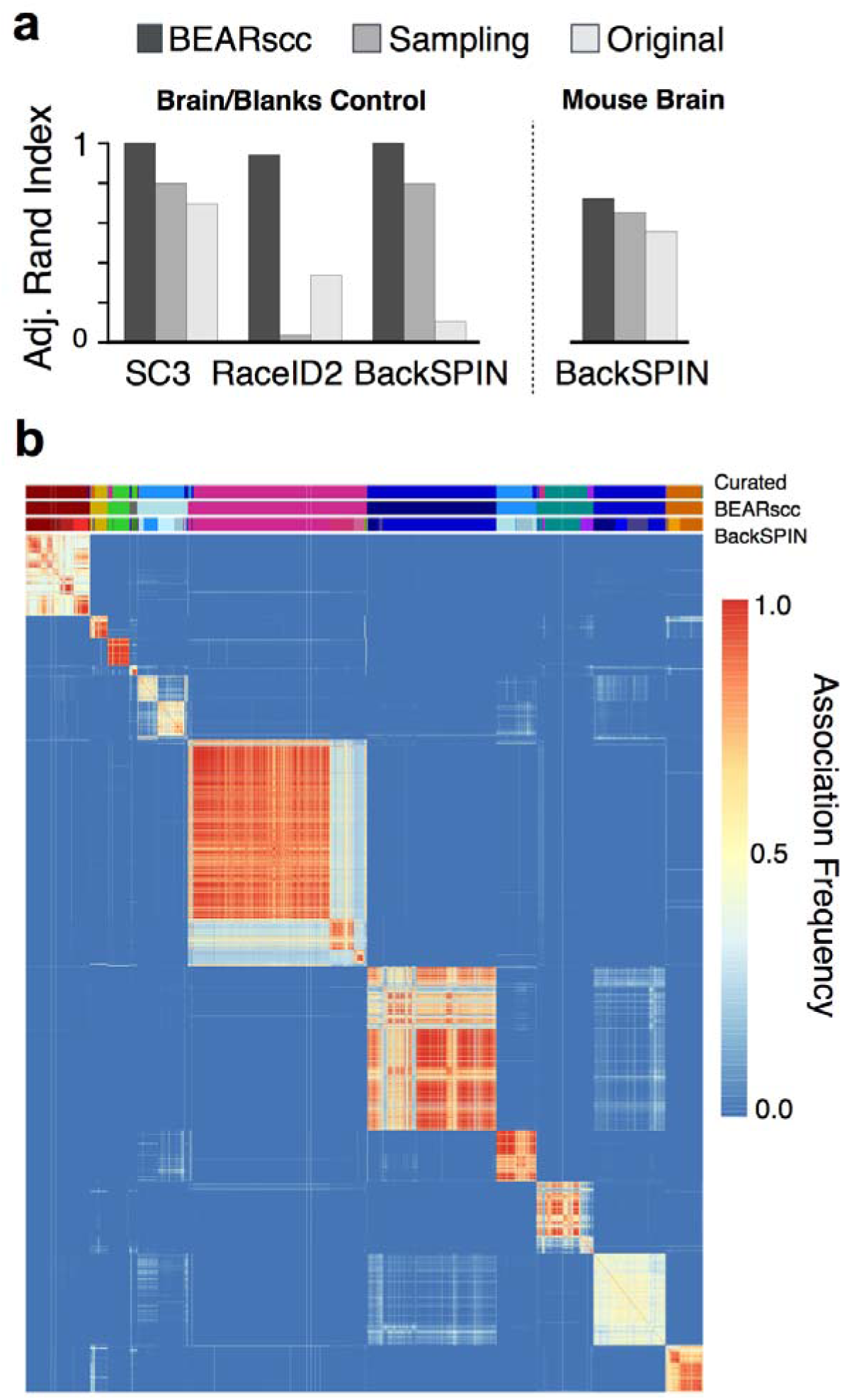
BEARscc improves clustering results and aids the interpretation of biological results. **a**, Comparison of clustering accuracy of control data (left) and murine brain data (right). Adjusted Rand index denotes agreement with the manually annotated grouping of samples (1: perfect, 0: no overlap). ‘BEARscc’ indicates that BEARscc was used to generate simulated technical replicates that were clustered using the algorithm indicated below the graph; ‘Sampling’ indicates that a sub-sampling approach (see text) was used before clustering with each algorithm; ‘Original’ indicates that the clustering algorithm was used alone. **b**, Example of a noise consensus matrix produced by BEARscc on data from murine brain cells (from Zeisel *et al.*) clustered with BackSPIN. Bars above heatmap show the manually curated clustering of cells (top), BEARscc consensus cluster (middle) and unsupervised BackSPIN clusters (bottom).

### BEARscc enhances analysis of published datasets

In order to assess the robustness of real biological data, we applied BEARscc to two previously published datasets from previously published scRNA-seq experiments. We first applied it to murine brain data (3005 cells) from Zeisel et al.^14^. Based on the ‘*score*’ statistic, BEARscc reduced the 24 clusters produced by BackSPIN (the algorithm used in the original publication) into 11 clusters which corresponded well with the manually curated cell types described in the original publication (Adjusted Rand Index 0.72 with BEARscc, and 0.55 for BackSPIN alone; Figure 2a (right), Figure 2b). Therefore, BEARscc provided an optimal grouping of cells without the effort of manual curation.

In a second evaluation, we re-analyzed murine intestinal data (291 cells) obtained by Grün *et al.*^13^, using BEARscc to generate simulated technical replicates and RaceID2 (as described in the original publication) for clustering. The ‘score’ metric from BEARscc indicated that 219 out of 291 cells were robustly classified in the original work. However, the two largest clusters - “cluster 1” and “cluster 2” - exhibited low *scores* (−0.07 and 0.20, respectively) compared to the other non-outlier clusters 3, 4 and 5 (Supplementary Figure 3a). The BEARscc noise consensus matrix reveals high variability in the clustering patterns of cells in clusters 1 and 2 (Supplementary Figure 3b). Grün et al. suggest that clusters 1 and 2 reflect closely related, undifferentiated cell types (“transit-amplifying” and “stem-like”, respectively). Expression patterns of genes characteristic of the two clusters were highly similar (Supplementary Figure 4a), compared to the expression differences between cluster 1 and the next-largest cluster (cluster 5) (Supplementary Figure 4b). Expression fold-changes between clusters 1 and 2 were reduced in technical replicates, falling below the significance threshold for many genes. BEARscc shows that many cells in clusters 1 and 2 cannot be reliably classified into one cluster or the other. The initially described sharp distinction between clusters 1 and 2 is therefore likely a result of technical variation, rather than a defining biological feature of these cells. ; Instead cells in clusters 1 and 2 seem to lie on a gradient of differentiation between two cellular states. More work will be needed to fully determine how the differentiation state of stem-like cells is reflected by their transcriptome. Nevertheless, this example demonstrates how BEARscc can help to improve the biological interpretation of scRNA data.

## Discussion

BEARscc addresses the challenges posed by intrinsically high technical variability in single-cell transcriptome sequencing experiments and enables the evaluation of single cell clustering results. Importantly, BEARscc is not a clustering algorithm in itself, but rather a tool to evaluate the results produced by any available clustering algorithm. To do so, it aggregates the information from exogenous control spike-ins across samples to create a model of both the expected variance of endogenous read counts as well as the likelihood of false-negative measurements (drop-outs). This represents a novel and alternative use of spike-in controls that is not subject to potential issues surrounding the use of spike-ins for per-sample normalisation. Our application of BEARscc to biological datasets demonstrates that BEARscc reduces over-clustering, is able to identify biologically relevant cell groups in an unsupervised way and provides additional insights for the interpretation of single-cell sequencing experiments.

We note that extreme batch effects with a multi-modal distribution of variance could skew BEARscc's noise model and lead to biased simulated replicates. We envision that future versions of BEARscc will attempt to detect and warn about such biases. Furthermore, while the drop-out model calculated by BEARscc is accurate for genes with an average expression of more than one count, there is still scope for improvement. Future work will focus on more precise models of drop-outs in the context of very low gene expression. As it stands, BEARscc enables users to identify the components of scRNA-seq clustering results that are robust to noise, thereby increasing confidence in those results for downstream analysis. Therefore, we recommend that future scRNA-seq analysis pipelines apply the best available clustering algorithm in conjunction with BEARscc in order to define the most biologically meaningful groups of cells for interpretation.

## METHODS

### Public data

Primary murine cortex and hippocampus single cell measurements for 3005 cells from Zeisel *et al.*^14^ were retrieved from the publicly available Linnarsson laboratory data repository (http://linnarssonlab.org/cortex/). Primary murine intestinal single cell measurements of 260 cells from Grün *et al.*^1^ were downloaded from the van Oudenaarden github repository (https://github.com/dgrun/RaceID).

### Implementation

All scripts necessary for implementing BEARscc are available from our bitbucket repository as an installable R package (https://bitbucket.org/bsblabludwig/bearscc).

### Algorithmic generation of simulated technical replicates

Simulated technical replicates were generated from the noise mixture-model and two drop-out models. For each gene, the count value of each sample is systematically transformed using the mixture-model, *Z*(*c*), and the drop-out injection, Pr(*X*= 0 | *Y* = *k*), and recovery, Pr (*Y*_*𝒿*_ = *y* | *Y*_*𝒿*_ = 0), distributions in order to generate simulated technical replicates as indicated by the following pseudocode:

~~~
FOR EACH gene, *𝒿*
       FOR EACH count, *c*
            IF *c* = 0
                   *n* (← SAMPLE one count, *y*, from Pr(*Y_𝒿_* = *y* | *X_𝒿_* = 0)
                   IF *n* = 0
                          *c* ← 0
                   ELSE
                          *c* ← SAMPLE one count from *Z*(*n*)
                   ENDIF
             ELSE
                   IF *c* ≤ *k*
                          *dropout* ← TRUE with probability, Pr(*X* = 0 | *Y* = *k*)
                          IF *dropout* = *TRUE*
                                 *c* ← 0
                          ELSE
                                 *c* ← SAMPLE one count from *Z*(*c*)
                          ENDIF
                   ELSE
                          *c* ← SAMPLE one count from *Z*(*c*)
                   ENDIF
            ENDIF
            RETURN *c*
      DONE
DONE
~~~

### Modelling noise from spike-ins

Technical variance was modelled by fitting a single parameter mixture model, *Z*(*c*) to th spike-ins’ observed count distributions. The noise model was fit independently for each spike-in transcript and subsequently regressed onto spike-in mean expression to define a generalized noise model. This was accomplished in three steps:

1. Define a mixture model composed of *poisson* and *negative binomial* random variables: *Z* ∼ (1 − *α*) * *Pois*(*μ*) + *α* * *NBin*(*μ*, *σ*)
2. Empirically fit the parameter, *α_i_*, in a spike-in specific mixture-model, *Z_i_*, to the observed distribution of counts for each ERCC spike-in transcript, *i*, where *μ_i_* and *σ_i_* are the observed mean and variance of the given spike-in. The parameter, *α_i_*, was chosen such that the error between the observed and mixture-model was minimized.
3. Generalize the mixture-model by regressing *α_i_* parameters and the observed variance *σ_i_* onto the observed spike-in mean expression, *μ_i_*. Thus the mixture model describing the noise observed in ERCC transcripts was defined solely by *μ*, which was treated as the count transformation parameter, *c*, in the generation of simulated technical replicates.

In step 2, a mixture model distribution is defined for each spike-in, *i*: *Z_i_*(*α_i_*, *μ_i_*, *σ_i_*) ∼ (1 − *α_i_*) * *Pois*(*μ_i_*) + *α_i_* * *NBin*(*μ_i_*, *σ_i_*). The distribution, *Z_i_*, is fit to the observed counts of the respective spike-in, where *α_i_* is an empirically fitted parameter, such that the *α_i_* minimizes the difference between the observed count distribution of the spike-in and the respective fitted model, *Z_i_*. Specifically, for each spike-in transcript, *μ_i_* and *σ_i_* were taken to be the mean and standard deviation, respectively, of the observed counts for spike-in transcript, *i*. Then, *α_i_* was computed by empirical parameter optimization; *α_i_* was taken to be the *α_i, j_* in the mixture-model, *Z_i, j_*(*α_i, j_*, *μ_i_*, *σ_i_*) ∼ (1 − *α_i, j_*) * *Pois*(*μ_i_*) + *α_i, j_* * *NBin*(*μ_i_*, *σ_i_*), found to have the least absolute total difference between the observed count density and the density of the fitted model, *Z_i_*. In the case of ties, the minimum *α_i, j_* was chosen.

In step 3, *α*(*c*) was then defined with a linear fit, *α_i_* = *a* * *log*2(*μ_i_*) + *b*. *σ*(*c*) was similarly defined, *log*2(*σ_i_*) = *a* * *log*2(*μ_i_*) + *b*. In this way, the observed distribution of counts in spike-in transcripts defined the single parameter mixture-model, *Z*(*c*), used to transform counts during generation of simulated technical replicates:

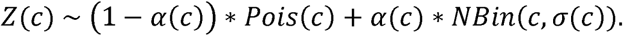

During technical replicate simulation, the parameter *c* was set to the observed count value, *a*, and the transformed count in the simulated replicate was determined by sampling a single value from *Z*(*c* = *a*).

### Inference of transcript drop-out distributions using spike-ins

A model of the drop-outs was developed in order to inform the permutation of zeros during noise injection. The observed zeros in spike-in transcripts as a function of actual transcript concentration and Bayes’ theorem were used to define two models: the ‘*drop-out injection distribution*’ and the ‘*drop-out recovery distribution*’.

The drop-out injection distribution was described by Pr(*X* = 0 | *Y* = *y*), where *X* is the distribution of observed counts and *Y* is the distribution of actual transcript counts; the density was computed by regressing the fraction of zeros observed in each sample, *D_i_*, for a given spike-in, *i*, onto the expected number spike-in molecules in the sample, *y_i_*, e.g. *D* = *a* * *y* + *b*. Then, *D* describes the density of zero-observations conditioned on actual transcript number, *y*, or Pr(*X* = 0 | *Y* = *y*). Notably, each gene was treated with an identical density distribution for drop-out injection.

In contrast, the density of the drop-out recovery distribution, Pr(*Y_𝒿_* = *y* | *X_𝒿_* = 0), is specific to each gene, *𝒿*, where *X_𝒿_* is the distribution of the observed counts and *Y_𝒿_* is the distribution of actual transcript counts for a given gene. The gene-specific drop-out recovery distribution was inferred from drop-out injection distribution using Bayes’ theorem and a prior. This was accomplished in 3 steps:

1. For the purpose of applying Bayes’ theorem, the gene-specific distribution, Pr(*X_𝒿_* = 0 | *Y_𝒿_* = *y*), was taken to be the the drop-out injection density for all genes, *𝒿*
2. The probability that a specific transcript count was present in the sample, Pr(*Y_𝒿_* = *y*), was a necessary, but empirically unknowable prior. Therefore, the prior was defined using the law of total probability, an assumption of uniformity, and the probability that a zero was observed in a given gene, Pr(*X_𝒿_* = 0). The probability, Pr(*X_𝒿_* = 0), was taken to be the fraction of observations that were zero for a given gene. This was done in order to better inform the density estimation of the gene-specific drop-out recovery distribution.
3. The drop-out recovery distribution density was then computed by applying Bayes’ theorem:

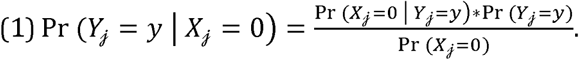

In the second step, the law of total probability, an assumption of uniformity, and the fraction of zero observations in a given gene were leveraged to define the prior, Pr(*Y_𝒿_* = *y*). First, a threshold of expected number of transcripts, *k* in *Y*, was chosen such that *k* was the maximum value for which the drop-out injection density was non-zero. Next, uniformity was assumed for all expected number of transcript values, *y* greater than zero and less than or equal to *k*; that is Pr(*Y_𝒿_* = *y*) was defined to be some constant probability, *n*. Furthermore, Pr(*Y_𝒿_* = *y*) was defined to be 0 for all *y* > *k*. In order to inform Pr(*Y_𝒿_* = *y*) empirically, Pr(*Y_𝒿_* = 0) and *n* were derived by imposing the law of total probability (2) and unity (3) yielding a system of equations:

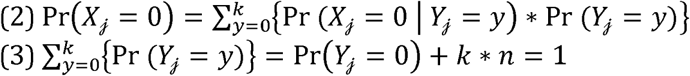

The probability that a zero is observed given there are no transcripts in the sample, Pr(*X_𝒿_* = 0 | *Y_𝒿_* = 0), was assumed to be 1. With the preceding assumption, solving for Pr(*Y_𝒿_* = 0) and *n* gives:

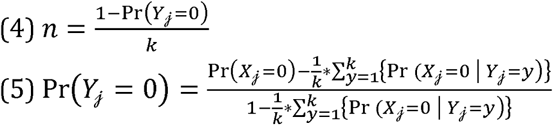

In this way, Pr(*Y_𝒿_* = *y*) was defined by (4) for *y* in *Y_𝒿_* less than or equal to *k* and greater than zero, and defined by (5) for *y* in *Y_𝒿_* equal to zero. For *y* in *Y_𝒿_* greater than *k*, the prior Pr(*Y_𝒿_* = *y*) was defined to be equal to zero.

In the third step, the previously computed prior, Pr(*Y_𝒿_* = *y*), the fraction of zero observations in a given gene, Pr(*X_𝒿_* = 0), and the drop-out injection distribution, Pr(*X_𝒿_* = 0 | *Y_𝒿_* = *y*), were utilized to estimate with Bayes’ theorem the density of the drop-out recovery distribution, Pr(*Y_𝒿_* = *y* | *X_𝒿_* = 0). During the generation of simulated technical replicates for zero observations and count observations less than or equal to *k*, values were sampled from the drop-out recovery and injection distributions as described in the pseudocode of the algorithm.

### Observing real technical noise

Brain whole tissue total RNA (Agilent Technologies, cat 540005) was diluted to 10pg aliquots and added to 1μL. cDNA conversion, library preparation, and sequencing were performed by the Wellcome Trust Center for Human Genomics Sequencing Core. Blank samples were identically prepared with nuclease free water. Samples were pipetted into 96-well plates and treated as single cells using Smartseq2 cDNA conversion as described by Picelli *et al*^16^ with minor modifications. The library was prepared using Fludigm's recommendations for Illumina NexteraXT at ¼ volume with minor modifcations, and sequenced on the Illumina HiSeq4000 platform. Raw reads were mapped to hg19 using STAR^17^. Exact position duplicates were removed, and features were counted using HTseq^17^.

### Clustering of counts data

BackSPIN, SC3 and RaceID2 were run according to algorithm-specific recommendations^13–15^. RaceID2 was allowed to identify cluster number under default parameters. For the brain and blanks control experiment data, RaceID2 was modified to skip normalization since scaled counts per million normalization had already been applied to the data set. The number of clusters, *k*, selected for SC3 clustering was determined empirically by selecting *k* with the optimal silhouette distribution across noise injected counts matrices.

### Computation of consensus matrix

100 simulated replicate matrices for *n* cells and *m* genes were clustered using the respective clustering algorithm (SC3, BackSPIN, RaceID2) as described above. Cluster labels were used to compute an *n* × *n* binary association matrix for each clustering. Each element of the association matrix represents a cell-cell interaction, where a value of 1 indicates that two cells share a cluster and a value of 0 indicates two cells do not share a cluster. An arithmetic mean was taken for each respective element across the resulting 100 association matrices to produce an *n* × *n* noise consensus matrix, where each element represents the fraction of noise injected counts matrices that, upon clustering, resulted in two cells sharing a cluster.

### Computation of BEARscc cluster metrics

To calculate cluster *stability*, the noise consensus matrix was subset to cells assigned to the cluster. The cluster *stability* was then calculated as the arithmetic mean of the upper triangle of the subset noise consensus matrix. To calculate cluster *promiscuity*, the rows of the noise consensus matrix were subset to cells assigned to the cluster and the columns are subset to the cells not assigned to the cluster. For clusters with as many or more cells assigned to them than not assigned, the *promiscuity* was defined as the arithmetic mean of the elements in the subset matrix. Otherwise, the columns were further subset to the same number of cells as were assigned to the cluster, where the cells outside of the cluster with the strongest mean association with cells inside the cluster are chosen. The *promiscuity* was defined as the arithmetic mean of the elements in this further subset matrix. Each cluster's *promiscuity* was subtracted from its *stability* to calculate cluster *score*.

### Computation of BEARscc cell metrics

To calculate a cell's *stability*, the arithmetic mean was taken of that cell's association frequencies with other cell's within the cluster. To calculate a cell's *promiscuity*, there were two cases. For cells in clusters with as many or more cells assigned to them than not assigned, the *promiscuity* was the arithmetic mean of that cell's association frequencies with all cells not assigned to the relevant cluster. For cells in clusters of size n, with fewer cells assigned to them than not assigned, the cell's *promiscuity* was the arithmetic mean with the *n* cells not assigned to the cluster with the highest association frequencies. Each cell's *promiscuity* was subtracted from its *stability* to calculate cell *score*.

### Estimation of cluster number *k*

In order to determine the cluster number, *k*, from the hierarchical clustering of the noise consensus, the resulting dendrogram was cut multiple times to form *N* clusterings with cluster numbers *k* = 1 to *k* = N clusters. The average *score* metric was computed for each clustering, and *k* was chosen by taking the *k* with the maximum average *score* metric. Evaluating all possible *k* from 1 to the number of cells in the experiment is computationally expensive and unlikely to be biologically meaningful. In this work, *N* was capped at 0.1 times the number of cells in the experiment: *N* = 10 for the brain and blanks control, *N* = 30 for the murine intestine experiment, and *N* = 300 for the murine brain data.

### Gene sampling

For comparison with BEARscc, 100 subsampling iteration matrices for *n* cells and *m* genes were generated by sampling one half of expressed genes and clustered using the respective clustering algorithm (SC3, BackSPIN, RaceID2). For each dataset, genes were excluded with less than 25 total raw counts across all samples in the cohort. The remaining genes formed the sample space. In each subsampling iteration, one half of the genes were sampled without replacement, and their expression across cells was used as the counts matrix. Identically to the computation of the BEARscc noise consensus matrix, cluster labels were used to compute an *n* × *n* binary association matrix for each clustering, and an arithmetic mean was taken for each respective element across the resulting 100 association matrices to produce an *n* × *n* subsampling consensus matrix. Identically to BEARscc analysis, the BEARscc *score* metric was used to determine cluster number *k*, and the resulting cluster labels for each dataset and algorithm were compared with BEARscc by computing the adjusted rand index for each with respect to the relevant ground truth.

### Author Contributions

D.T.S conceived and implemented the computational approach under the supervision of B.S.-B. M.W., R.O., and X.L. designed the initial experimental study that that led to the development of the presented approach and included the sequencing of brain and blanks samples. B.S.-B. and D.T.S. prepared the manuscript, with contributions from all authors.

### Acknowledgements

We thank Mary Muers, Andy Roth and Chris Ponting for careful reading of the manuscript, and Rory Bowden, Amy Trebes and the High-Throughput Genomics team at the Wellcome Trust Centre for Human Genetics for assistance with sequencing. All authors acknowledge support from Ludwig Cancer Research. DTS was supported by Nuffield Department of Clinical Medicine and the Clarendon Fund. MJW was supported by Cancer Research UK. MJW and RPO received funding from the NIHR Biomedical Research Centre. RPO received funding from Oxford Health Services Research Committee and Oxford University Clinical Academic Graduate School.

## Supplementary Material

**Supplementary Figure 1.**
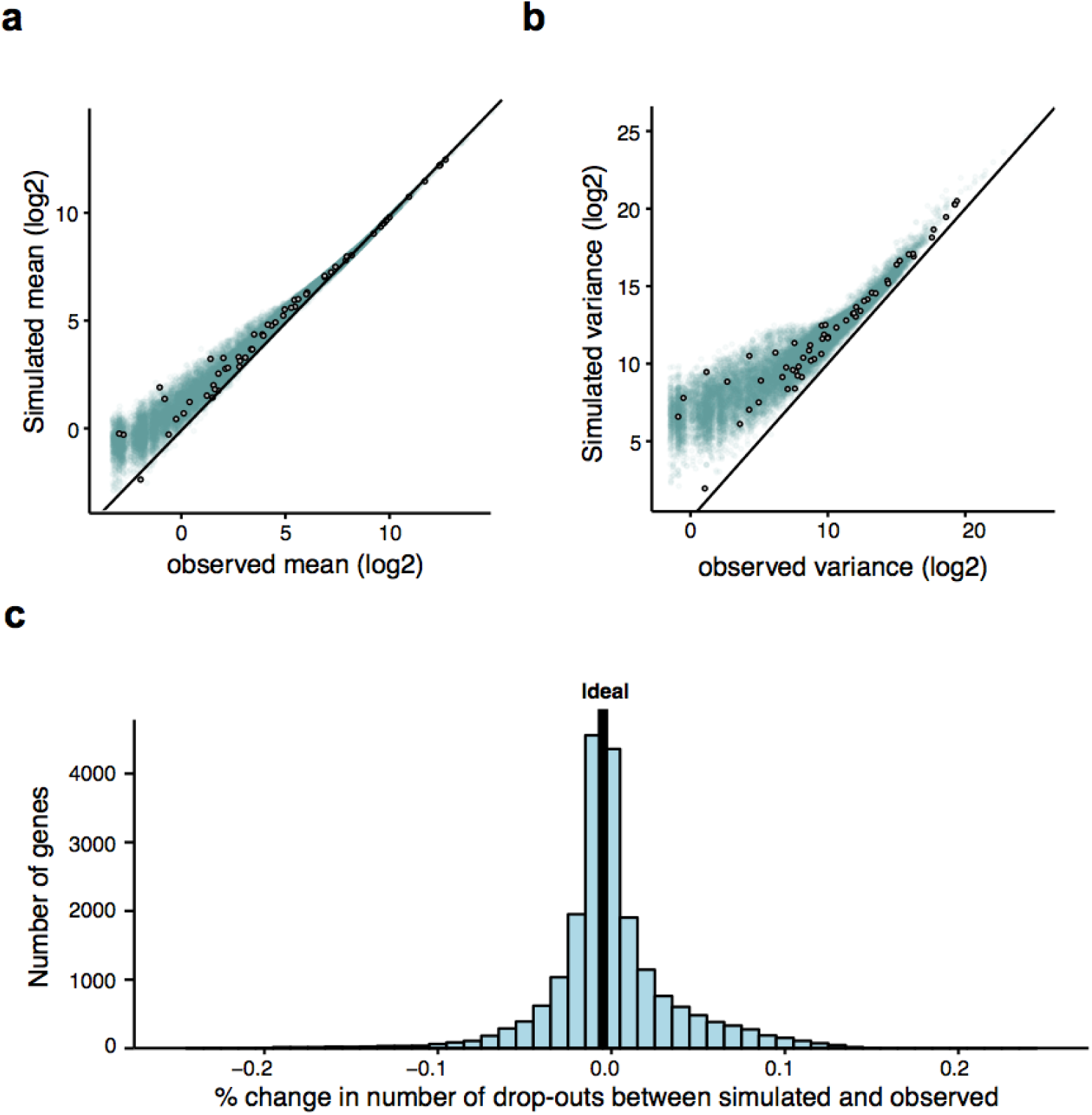
BEARscc accurately models technical variability. Scatterplots of observed vs simulated mean expression **(a)** and variance in expression **(b)**, based on data from brain RNA control experiment. ERCC spike-in values are circled in black, human genes are shown in blue. c, Difference between simulated and observed drop-out frequency across genes.

**Supplementary Figure 2.**
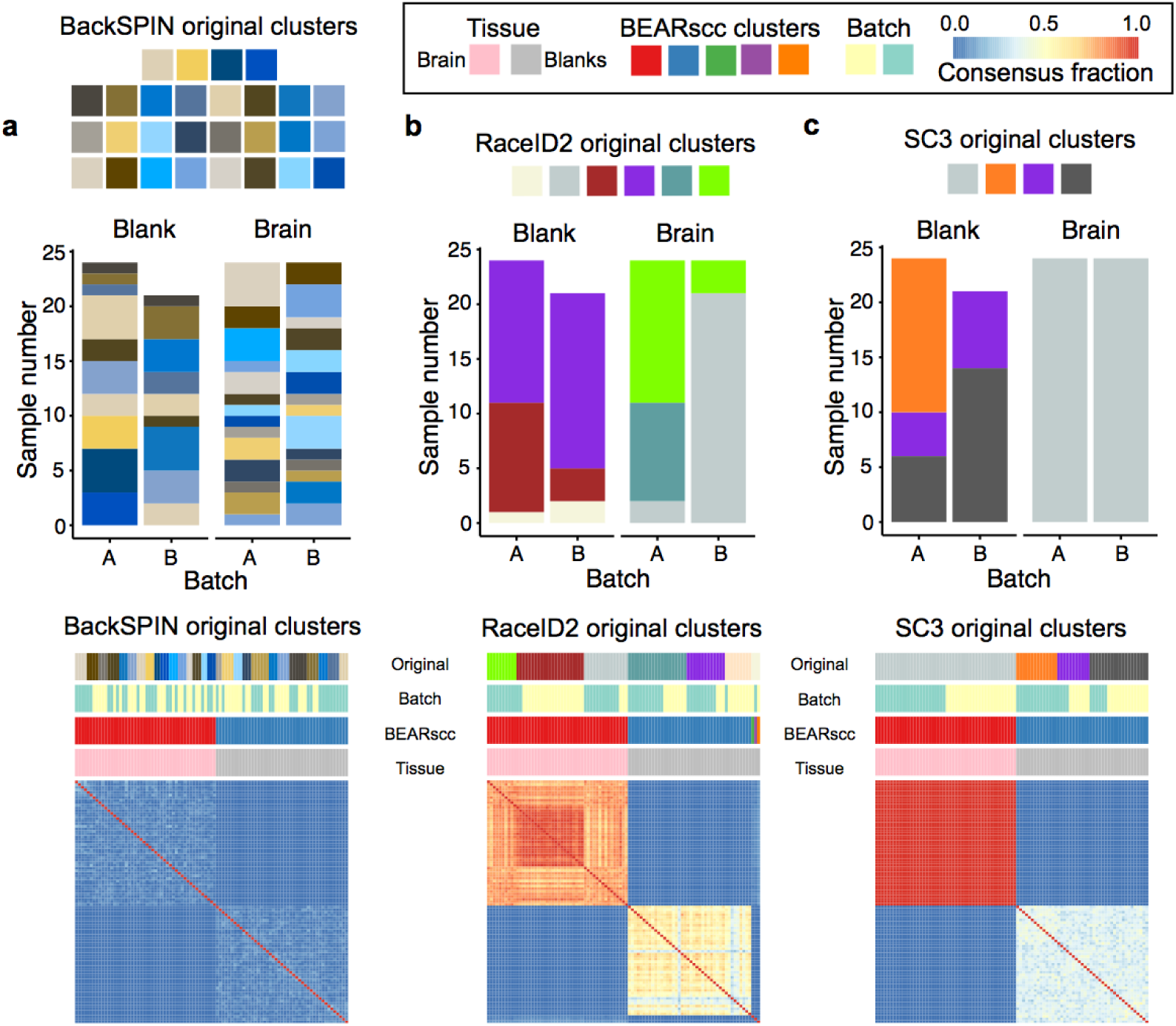
BEARscc applied to the brain-and-blanks control experiment in combination with BackSPIN **(a)**, RaceID2 **(b)** and SC3 **(c)**. Top: bar graphs showing how the clusters generated by using each clustering algorithm alone (‘original clusters’) relate to sample type (brain or blank) and batch (A or B). RaceID2 and SC3 clusters are visibly confounded by batch. Bottom: for BEARscc applied with each algorithm, the noise consensus matrix is shown. The bars above the matrix show (from top): original clusters with algorithm alone, the batch, clusters derived after application of BEARscc, and the sample type.

**Supplementary Figure 3.**
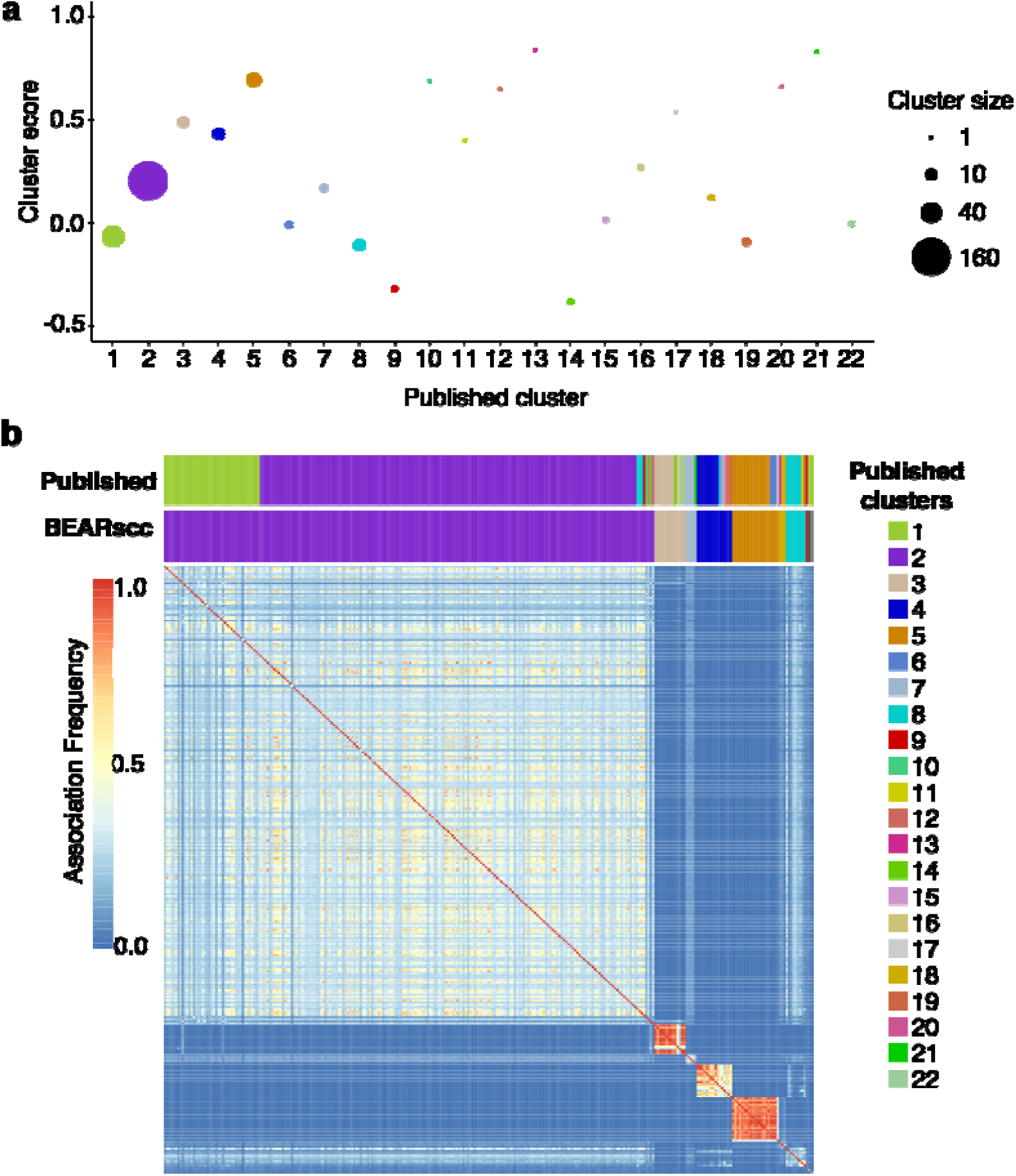
BEARscc identifies robust clusters in data from murine intestinal cells. **A**, Cluster scores for “main” clusters (1-5) and outlier clusters (6-22). Circle size reflects number of cells per cluster. Colors are the same as in subfigure b. **b**, BEARscc noise consensus matrix for murine intestinal cells clustered with RaceID2. Above heatmap: published clusters (top) and noise consensus clustering (bottom, colors indicate closest match in the published clustering).

**Supplementary Figure 4.**
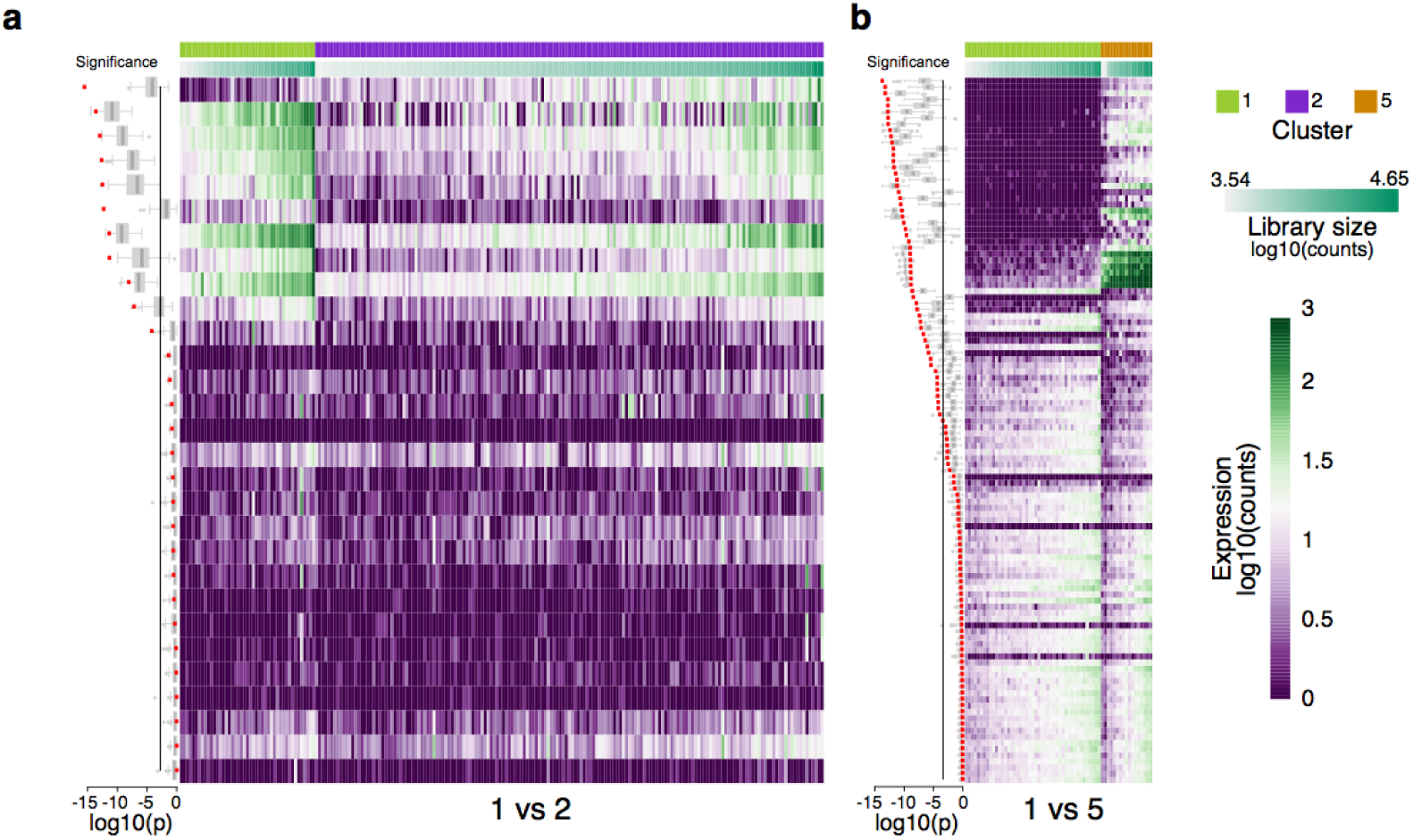
BEARscc correctly detects that separation of “stem-like” cell clusters 1 and 2 is based on weak expression differences. **(a)** Heatmap of expression of genes characteristic of clusters 1 and 2 (as described in the original manuscript), and **(b)** clusters 1 and 5. Columns in each heatmap are ordered by library size per cell, rows sorted by significance of expression fold-change between clusters. Boxplots on the left denote the significance of difference in expression between the two clusters (Wilcoxon rank-sum test). Red denotes the observed values, and simulated technical replicates are shown in gray. Black solid vertical line denotes Bonferroni-corrected significance threshold.

## References

1. Grün, D. et al. Single-cell messenger RNA sequencing reveals rare intestinal cell types. Nature 525, 251–255 (2015).

2. Tirosh, I. et al. Dissecting the multicellular ecosystem of metastatic melanoma by single-cell RNA-seq. Science 352, 189–196 (2016).

3. Grün, D., Kester, L. & van Oudenaarden, A. Validation of noise models for single-cell transcriptomics. Nat Methods 11, 637–640 (2014).

4. Kim, J. K. et al. Characterizing noise structure in single-cell RNA-seq distinguishes genuine from technical stochastic allelic expression. Nat Commun 6, 8687 (2015).

5. Hicks, S. C., Teng, M. & Irizarry, R. A. On the widespread and critical impact of systematic bias and batch effects in single-cell RNA-Seq data. BioRxiv (2015). doi:10.1101/025528

6. Jiang, L. et al. Synthetic spike-in standards for RNA-seq experiments. Genome Res 21, 1543–1551 (2011).

7. Vallejos, C. A., Marioni, J. C. & Richardson, S. BASiCS: Bayesian Analysis of Single-Cell Sequencing Data. PLoS Comput Biol 11, e1004333–18 (2015).

8. Brennecke, P. et al. Accounting for technical noise in single-cell RNA-seq experiments. Nat Methods 10, 1093–1095 (2013).

9. Wagner, A., Regev, A. & Yosef, N. Revealing the vectors of cellular identity with single-cell genomics. Nat Biotechnol 34, 1145–1160 (2016).

10. Rousseeuw, P. J. Silhouettes: A graphical aid to the interpretation and validation of cluster analysis. J Comput Appl Math 20, 53–65 (1987).

11. Tibshirani, R., Walther, G. & Hastie, T. Estimating the number of clusters in a data set via the gap statistic. J Royal Statistical Soc B 63, 411–423 (2001).

12. The External RNA Controls Consortium: a progress report. 2, 731–734 (2005).

13. Grün, D. et al. De Novo Prediction of Stem Cell Identity using Single-Cell Transcriptome Data. Stem Cell 19, 266–277 (2016).

14. Zeisel, A. et al. Cell types in the mouse cortex and hippocampus revealed by single-cell RNA-seq. Science 347, 1138–1142 (2015).

15. Kiselev, V. Y. et al. SC3 - consensus clustering of single-cell RNA-Seq data. BioRxiv (2016). doi:10.1101/036558

16. Picelli, S. et al. Full-length RNA-seq from single cells using Smart-seq2. Nat Protoc 9, 171–181 (2014).

17. Dobin, A. et al. STAR: ultrafast universal RNA-seq aligner. Bioinformatics 29, 15–21 (2012).

